# Machine Learning Gene Signature to Metastatic ccRCC based on ceRNA Network

**DOI:** 10.1101/2023.07.31.551358

**Authors:** Epitácio Farias, Patrick Terrematte, Beatriz Stransky

## Abstract

Renal carcinoma is a pathology of silent and multifactorial development characterized by a high rate of metastases in patients. After several studies have elucidated the activity of coding genes in the metastatic progression of renal carcinoma, new studies seek to evaluate the association of non-coding genes, such as competitive endogenous RNA (ceRNA). Thus, this study aims to build a gene signature for clear cell renal cell carcinoma (ccRCC) associated with metastatic development from a ceRNA network and to analyze the probable biological functions performed by the participants of the signature. Using ccRCC data from The Cancer Genome Atlas (TCGA), we constructed the ceRNA network with the differentially expressed genes, assembled nine gene signatures from eight feature selection techniques, and analyzed the evaluation metrics of the classification models in the benchmarking process. With the signature, we performed somatic and copy number alteration analysis, survival and metastatic progression risk analysis, and functional annotation analysis. In this study, we present an 11-gene signature (SNHG15, AF117829.1, hsa-miR-130a-3p, hsa-mir-381-3p, BTBD11, INSR, HECW2, RFLNB, PTTG1, HMMR, RASD1). Validation using the external dataset of the International Cancer Genome Consortium (ICGC-RECA) made it possible to assess the generalization of the signature, which showed an Area Under Curve of 81.5%. The genomic analysis identified the signature participants on chromosomes with highly mutated regions (G-index > 2). The hsa-miR-130a-3p, AF117829.1, hsa-miR-381-3p, and PTTG1 had a significant relationship between expression and patient survival, and the first two had a significant association with metastatic development. In addition, functional annotation resulted in relevant pathways for tumor development, such as PI3K/AKT, TNF, FoxO, RNA polymerase two transcription regulation, and cell control. Finally, by analyzing the connections of the signature genes within the ceRNA network in conjunction with studies in the literature, it was possible to obtain an overview of their activities within the ccRCC. Therefore, this gene signature identified new coding and non-coding genes and could act as potential biomarkers for a better understanding of renal carcinoma and in the development of future treatments in the clinical area.

## 1. Introduction

Renal cancer is a group of neoplasms originating in the renal tissues, classified by the cell type or histologic characteristics, such as Clear Cell Renal Cell Carcinoma (ccRCC), Papillary Renal Carcinoma (pRCC), and Chromophobe Renal Carcinoma (chRCC)[1–3]. Due to the silent characteristic of this disease [4], the diagnosis at the metastatic state occurs in approximately 30% of ccRCC patients [5, 6].

In a previous study with a cohort of 537 ccRCC patients, The Cancer Genome Atlas (TCGA) consortium [7], characterized significant alterations in ccRCC, such as mutation in VHL, PBRM1, SETD2, BAP1 genes, the deletion of de arm q of chromosome 3, and a cluster organization with messenger RNA (mRNA) and microRNA (miRNA), representing an essential component in ccRCC regulation. Further studies begin to reveal an important role of the non-coding RNAs (ncRNAs) represent the class of RNAs that portray approximately 80% of the transcriptome [8–10].

The function of lncRNAs is associated with the location of action or their interactions with DNA, proteins, or other RNAs [9–13]. The lncRNAs can act during all the transcriptional processes, as the pre and post-transcriptional processes, as a: (i) decoy or “sponge” modulating the effector of their targets; (ii) guide to the enzymes modifiers of histones or chromatin; (iii) respond to various stimuli [14, 15]. The ligation of the lncRNA with the miRNA affects their targets, characterizing an endogenous competition between the lncRNA and the mRNA target of the miRNA [9, 10].

Based on this widespread interaction network, it was proposed the “Competing Endogenous RNA” (ceRNA) hypothesis, based on the idea of a communication existence between miRNAs, mediated by the miRNAs recognition elements (MREs), with mRNA, lncRNA, and other ncRNAs[16]. Alteration in the ceRNA networks is observed in cancer and other pathologies, associating them with biomarkers of prognosis to metastasis and alternative clinical outcomes, therapeutics targets, where they can act as a tumor suppressor or oncogenes [10, 17–20].

Studies about RNA expression generate a large and complex amount of data, and the conduction of an analysis integrating this data with clinical information could enable a pattern extraction to enrich the understanding by machine learning (ML) techniques [21, 22]. Among the vast applications of ML, the methods related to classification and prediction became the most used approach in health field research [23]. However, the lack of feature selection associated with the outcome variable could influence the performance of the algorithms [24]. The feature selection represents the analysis and selection of variables, evaluating their impact on the outcome, removing the irrelevant variables, and making them more consistent and relevant to the model construction [25].

This study aims to construct a ceRNA network and a gene signature based on the feature selection algorithms, to classify the metastatic profile of ccRCC patients. The best-performing gene signature achievement used majority voting between four Recursive Feature Elimination (RFE) approaches. More specifically, the RFE is a wrapper-based method to select the classifiers interactively, initially using all the variables, and for each interaction, one variable is removed based on the score of importance associated [26]. The flowchart shown in Figure 1 displays a summarized view of the discovery process for the novel RFE gene signature of ccRCC.

**Figure 1:**
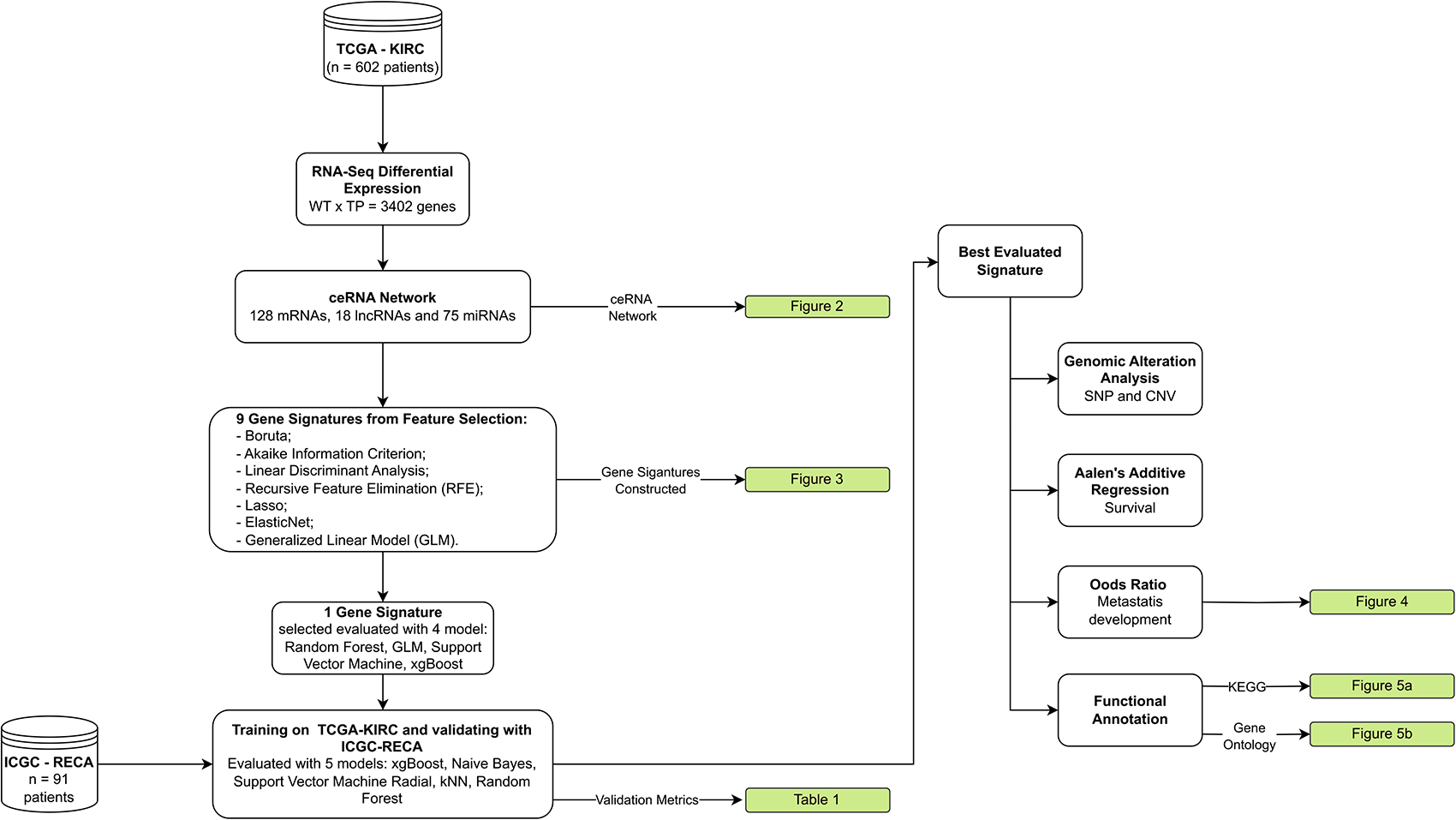
Flowchart of the current study to obtain a gene signature based on the Recursive Feature Elimination (RFE) approach. The datasets are indicated by the cylindric shape, the white rectangles represent the steps of the study, and the green rectangles represent the resulting Figures and tables. TCGA-KIRC and ICGC-RECA are the ccRCC datasets.

## 2. Materials and Methods

### 2.1. Data

This is a data-driven study based on the RNA-seq dataset and clinical dataset from the TCGA-KIRC project (n = 602), downloaded from Genomic Data Commons (https://portal.gdc.cancer.gov/) [7] and UCSC Xenabrowser (https://xena.ucsc.edu/). For external validation, we used the dataset of ccRCC (n = 91 patients) from the International Cancer Genome Consortium (ICGC-RECA) [27].

### 2.2. ceRNA Network construction

The ceRNA network was constructed from the differentially expressed genes mRNAs and ncRNAs, focusing on the relation lncRNA-miRNA-mRNA. The differential expression analysis was made between the normal tissues (n = 72) and tumor tissues (n = 530) from the TCGA-KIRC cohort, with the “DESeq2” using the absolute |log-fold change (LFC)| > 2 and a *p-value* adjusted (FDR) < 0.01.

With the differentially expressed genes, was used the R package “GDCRNATools” [28] associated with the starBase [29], a database focused on the decodification of the iterations networks through numerous RBPs and RNAs. The pair selection follows the statistical analyzes: (i) hypergeometric test; (ii) Pearson correlation coefficient; (iii) regulatory similarity. This analysis used a threshold of 0.1 for the Pearson correlation and hypergeometric test and 0 for the regulatory similarity, and Cytoscape software [30] to visualize the ceRNA network.

### 2.3. Dataset Construction, Feature Selection, and Gene Signature Construction

The signature construction used the genes participating in the ceRNA network inspired by the methodology of [31], where new gene signatures were produced using the techniques in Table S1 and used the OmicSelector R package [32].

Within the expression dataset from the TCGA-KIRC (n = 602) was observed a missing metastatic classification in 30 patients, causing their remotion, and due to the unbalanced characteristic from the metastasis classification of presence (M1) or absence (M0), was performed a propensity matching score balance, maintaining 190 patients, 95 from each class.

This new dataset was split randomly into three new datasets, following the rate of 60% for training (n = 114), 20% for the test (n = 38), and 20% for validation (n = 38). For the signature construction process, we used the feature selection techniques: Recursive Feature Elimination (RFE) and two iterated versions, Boruta, Generalized Linear Model (GLM), Akaike Information Criterion (AIC), Linear Discriminant Analysis (LDA), Lasso and ElasticNet.

To improve the construction of the signature and optimize computational efficiency, we performed hyperparameters adjustments to the feature selection. The RFE techniques used cross-validation with ten folds, using a window frame of 50 genes in each iteration, and iterated RFE versions used a window frame of ten genes for the signature.

With the nine signatures constructed, was performed a 1° benchmarking to select the signature with the best metrics for metastatic classification using the datasets for test and validation. To the 1° benchmark was used the models: Random Forest (rf), Generalized Linear Model (GLM), eXtreme Gradient Boosting (xgbTree) e Support Vector Machine with a Radial Kernel (svmRadial), performed ten times to seek the best parameter adjustment for each of them. The metrics to evaluate this benchmark are accuracy, specificity, sensitivity, and Youden’s statistics.

To evaluate the signature generalization, the external dataset from the ICGC-RECA project (n = 91) was used with the mlr3verse package [33] to perform the 2° benchmark, applying the following classification techniques: random forest, naive Bayes, kNN, svmradial, and XGBoost. The evaluation metrics were accuracy, balanced accuracy, the Brier score, and the AUC. The validation process used the TCGA-KIRC for training and the ICGC-RECA for testing.

### 2.4. Somatic and Copy Number Alteration Analysis

The somatic alterations analysis was conducted with the Mutation Annotation Format (MAF) datafile, using the R package, Maftools [34], extracting information about (a) type of variations; (b) variation classification; (c) the labels of those single nucleotide variations; (d) the variations quantity by sample and (e) the top 10 genes altered.

The copy number variation analysis requires the construction of the GISTIC file. The Genomic Identification of Significant Targets in Cancer (GISTIC) pipeline [35] resulted in information about amplification and deletions within the data, analyzed by the Maftools R package to extract the regions of the genome and their alterations.

### 2.5. Risk Analysis

The performance of risk analysis allows assessing the relationship between the gene signature with the metastatic development and the survival status of the patients, observing their expression level.

With the survival [36] and finalfit [37] R packages, we executed Aalen’s additive regression and Odd’s ratio analysis, respectively. Aalen’s regression acts as a complementary, or alternative, form for the Cox model, where the covariables association and their effects [38] on the survival status of the patients are obtained. The Odd’s ratio quantifies the strength of association between two events [39], the presence or absence of metastasis.

### 2.6. Functional Annotation Analysis

The identification of the pathways enriched by the genes of the signature was performed against the Kyoto Encyclopedia of Genes and Genomes (KEGG) [40] and the Gene Ontology [41], focusing on the gene association to biological processes and molecular functions. Using the clusterProfiler R package [42] and the mirPath platform [43] for functional characterization of miRNAs from the signature.

## 3. Results

### 3.1. ceRNA Network

To construct our ceRNA network, we used the differentially expressed (DE) genes of the TCGA-KIRC (n = 602) project. This analysis resulted in 2,842 mRNAs, 132 miRNAs, and 271 lncRNAs DE, based on the thresholds of |log2FC| > 2 and p-value adjusted for FDR < 0,01.

With those DE genes, we constructed the ceRNA network based on the thresholds of 0.1 for the hypergeometric test and Pearson correlation, and the similarity of regulation different from 0, resulting in a network with 18 lncRNAs, 75 miRNAs and 128 mRNAs (Figure 2).

**Figure 2:**
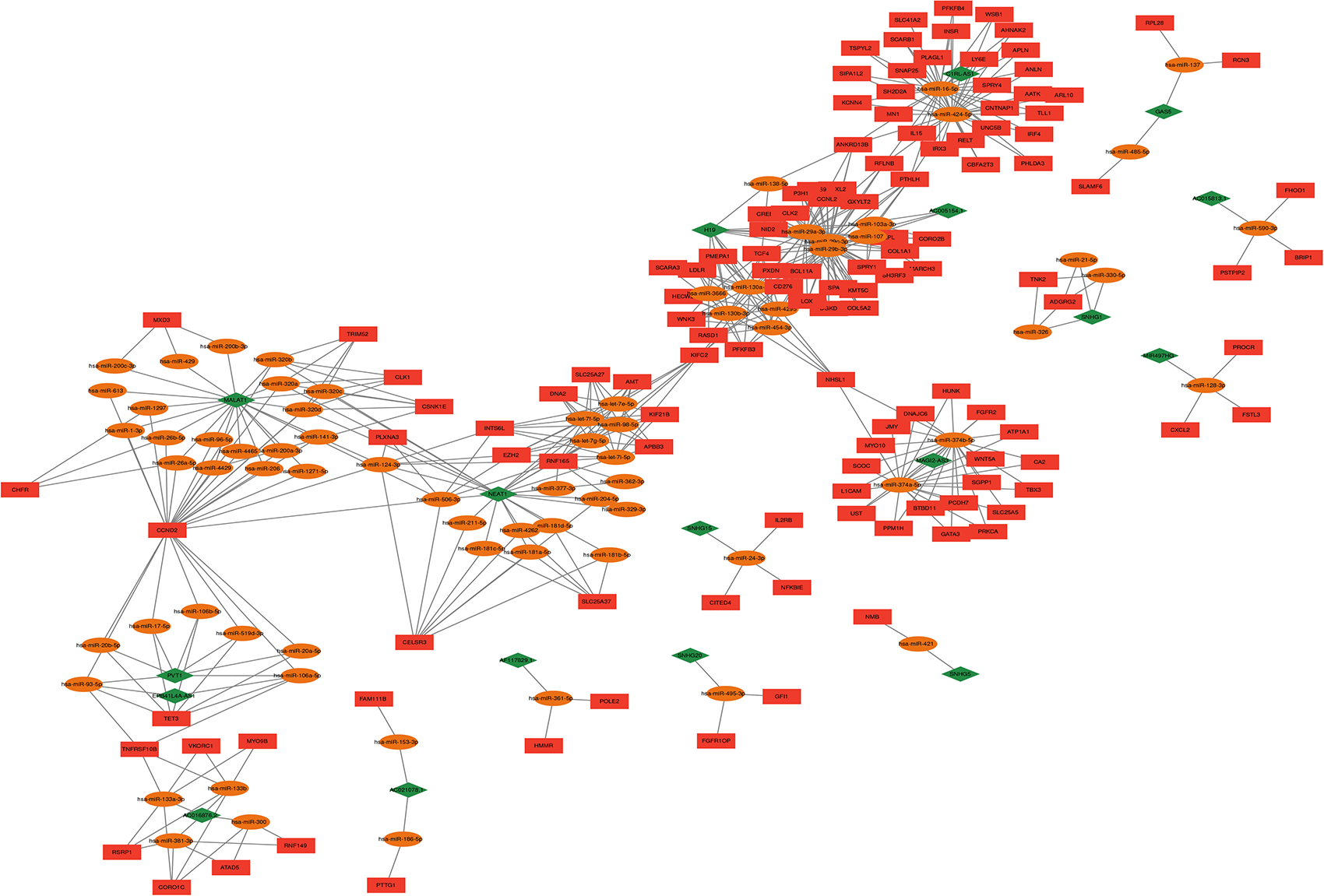
The ceRNA network constructed based on the difierentially expressed (DE) genes in the ccRCC patients. It’s observed a cluster conformation, were exist regions highly connected, and regions slightly connected. The red rectangles represent the messenger RNAs (mRNAs), the orange elipses represents the micro RNAs (miRNAs), and the green losang represents the long non-coding RNAs (lncRNA). The network is composed by 18 lncRNAs, 75 miRNAs, and 128 mRNAs.

### 3.2. Feature Selection

With the expression data from the 221 genes participating in the ceRNA network and the metastatic classification from the 192 patients, after the balance performance, the training process for the feature selection and the construction of 9 signatures were performed (Figure 3). Among the feature selection techniques, only the ***stepAIC*** did not converge, and the curves from RFE show an accuracy of 76.30% and a Kappa coefficient of 0.5663.

**Figure 3:**
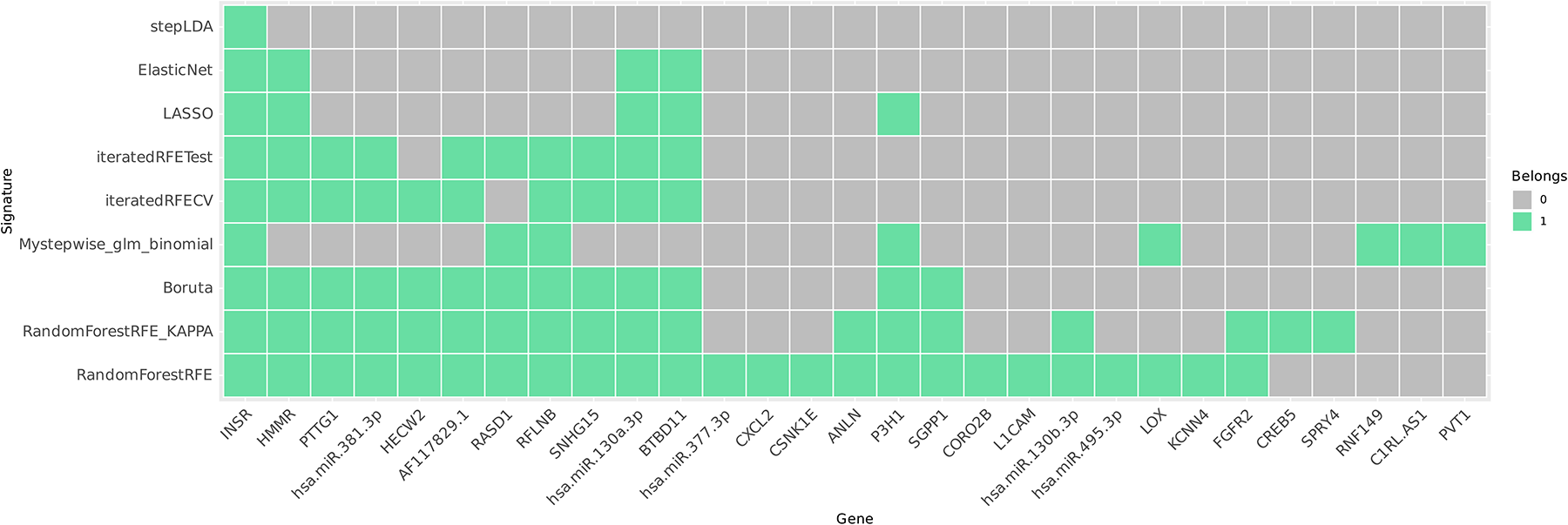
Heatplot with the 29 unique genes reported by the 9 gene signature constructed. In the Y-axis are the methods applied to the signature construction, and in the X-axis are the genes listed. The green square represent the presence of the gene as resulted by the method

After the stepAIC remotion, we performed the first benchmark, where the *xgbTree* presented the best result, with an accuracy of 80% during the training and 60% for the test, and 68.3% in validation. To select the best signatures, we applied Youden’s statistics, resulting in the best four signatures. As observed, the four signatures shared some genes, and by majority voting, the final signature was constructed (Equation 1).

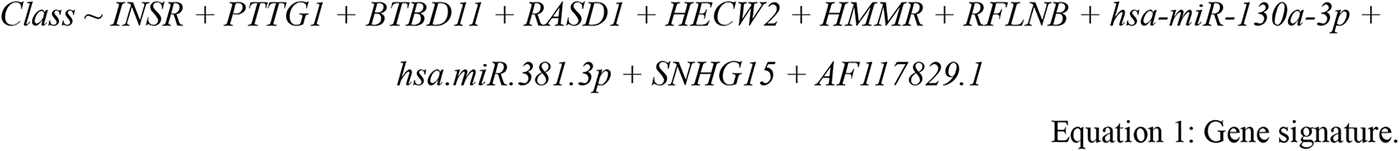

With the signature constructed, a second benchmark (Table 1) was performed, using the ICGC-RECA project as a test dataset, and observed accuracy and balanced accuracy of 72% for both, an AUC of 81.5%, and a Brier Score of 0.1955.

**Table 1:** Metrics evaluated for validation with an external dataset.

| Method | Accuracy | Balanced Accuracy | AUC | Brier Score |
| --- | --- | --- | --- | --- |
| RandomForest | 72.2% | 72.2% | 81.48% | 0.1955442 |
| SVM | 50% | 50% | 66.67% | 0.2500714 |
| xgBoost | 61.1% | 61.1% | 62.34% | 0.2343498 |
| kNN | 50% | 50% | 61.72% | 0.4817816 |
| Naive Bayes | 50% | 50% | 54.32% | 0.5000000 |

### 3.3. Integrative Analysis From The Transcriptional Signature Components

#### 3.3.1. Genomic Alteration Analysis

Performing a genome-level alteration analysis enables us to evaluate their impact on the gene product. These alterations can include changes in the genetic structure, disruptions in protein synthesis, or variations in the quantity of the gene product. To conduct this analysis, we used the maftools package to investigate single nucleotide polymorphisms (SNPs) and copy number variations (CNVs) within the genome of the TCGA-KIRC cohort.

Among the data, the missense mutation is predominant from the single nucleotide polymorphism type, with approximately 44 variants per sample. The most common SNP was the cytosine and thymine transversion.

As the focus is on the gene signature, ten samples showed mutations in signature coding genes (Figure S1), where the missense was registered at the genes HECW2, BTBD11, INSR, and PTTG1, the frameshift deletion was registered in BTBD11, and the multi-hit mutations in HECW2. However, the HMMR, RASD1, and RFLNB have not presented any variation.

The copy number variation analysis shows the chromosomes 1,4,5,6,7,12,17,18, and 20 with a large amount and frequency of alterations between the samples. As we searched the chromosome location of our gene signature in the *National Center of Biotechnology Information*, we observed that their localization was in the chromosomes highly altered but not in the regions significantly modified.

#### 3.3.2. Risk Analysis

As we construct the risk analysis associating the gene signature expression with the ccRCC progression, Aalen’s additive regression shows a significant relationship between some genes from the gene signature with patient survival, such as (i) *AF1117829.1* (p-valor = 0,0001627), (ii) *hsa-miR-130a-3p* (p-valor = 0,016), (iii) *hsa.miR.381.3p* (p-valor = 0,027) e (iv) *PTTG1* (p-valor = 0,020).

When observing the behavior of the signature genes with the development of metastasis (Figure 4), the miRNA hsa-miR-130a-3p and the lncRNA AF117829.1 were the only ones that had a significant association, p-value = 0.011, and p-value = 0.029, respectively.

**Figure 4:**
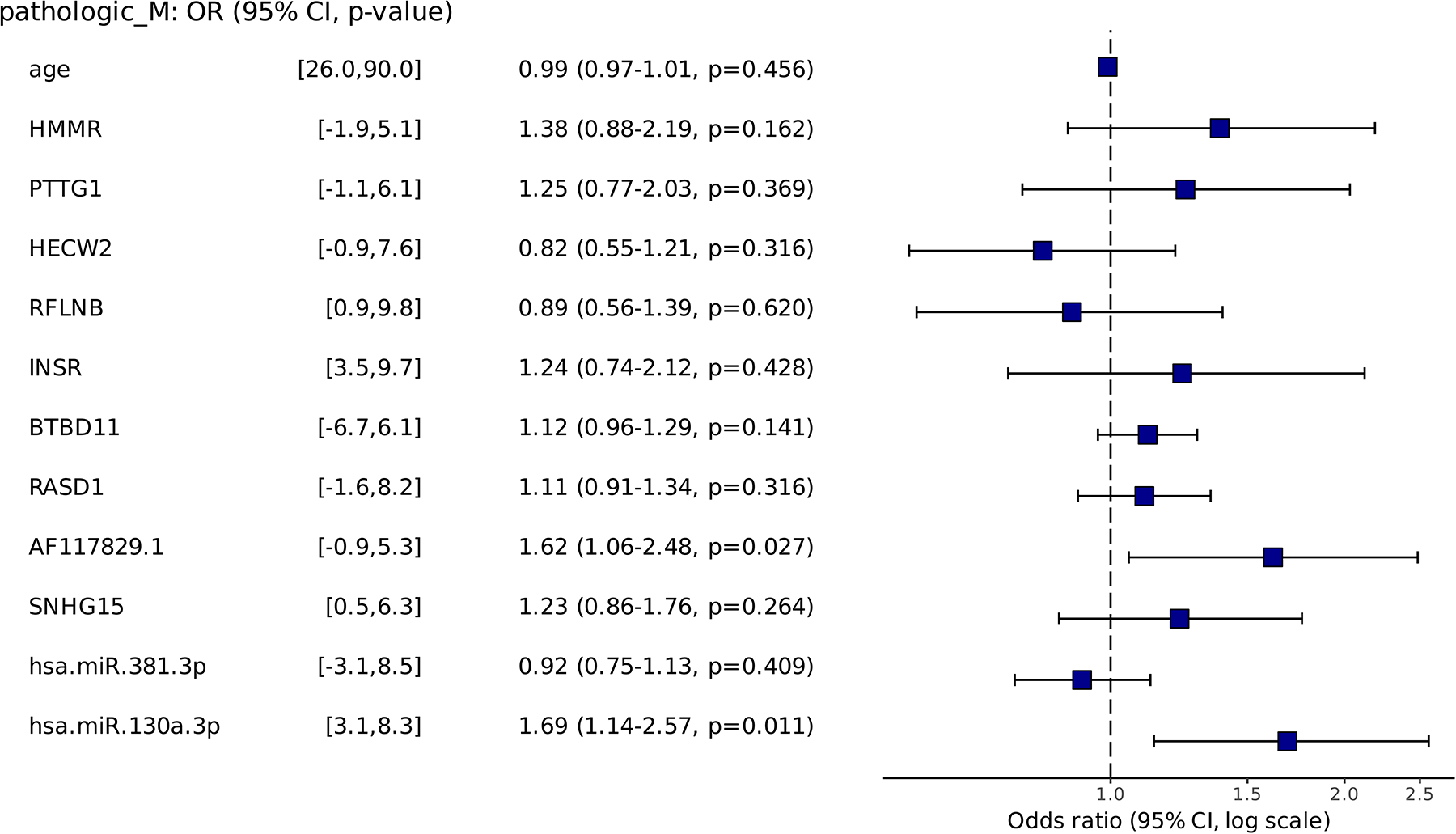
Odds ratio of each gene in the signature regarding metastatic development and 95% confidence interval. The miRNA hsa-miR-130a-3p and the lncRNA AF117829.1 were the only ones significantly associated (p-value < 0.05).

#### 3.3.3. Functional Annotation Analysis

During the annotation of KEGG pathways based on the coding genes from the signature, we observed an association between several biological pathways, like longevity regulation, and aldosterone-regulated sodium reabsorption, with a p-value <0.05 (Figure 5a). When evaluating the targets from the miRNAs in our signature and the biological pathways related to them, it resulted in well-known oncology-related biological pathways, like the PI3K-AKT signaling pathway, p53 signaling pathway, the transforming growth factor-beta (TGF-beta) signaling pathway, renal cancer, and HIF-alfa pathway (Figure S2a).

**Figure 5:**
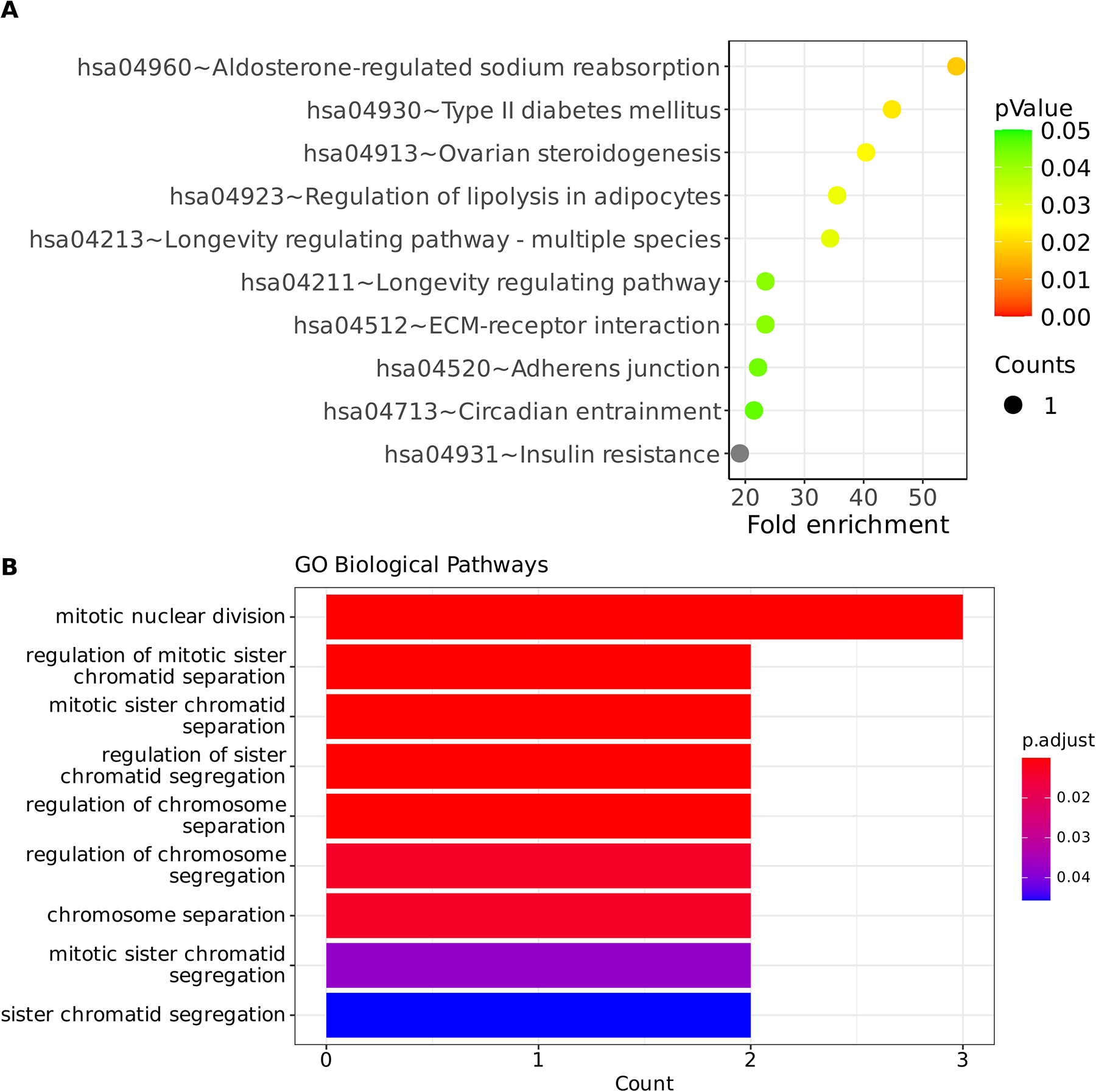
Functional annotation from (a) KEGG for the seven encoding genes and from (b) Gene Ontology for the target mRNAs of the two miRNAs involved in the signature. In both cases, the Y-axis represents the annotated pathways for their respective input data, while the X-axis for figure (a) represents the membership relationship between the signature mRNAs and the total genes in the pathway, and for figure (b) it represents the number of genes from signature in the pathway.

The biological processes annotated were associated with cellular division regulation, like chromatid sister separation and chromosome segregation (Figure 5b). The pathways annotated by miRNA targets were also related to the cellular division process. However, other pathways were listed, like the signal transduction pathway, growth factors, and DNA polymerase I regulation, both analyses with a p-value < 0.05 (Figure S2b).

### 3.4. Gene Signature and ceRNA network

As the signature construction was made upon the genes from the ceRNA network, searching their location and the first neighbors could improve the knowledge about the gene functions and their metastatic consequences in the ccRCC environment.

The ceRNA network had a cluster organization, and the gene signature location showed the presence of genes in areas with cluster distinct, like cluster 1, or with a high density of connections, like cluster 2, or even areas with the presence of only one gene like the cluster 3,4,5, and 6, Table 2 present the genes from signature and their first ligands within the ceRNA network.

## 4. Discussion

### 4.1. Gene Signature

The nine feature selection methods resulted in the training of signatures for the metastatic classification of ccRCC. Analyzing the learning curves from RFE, the Kappa coefficient between the range of 0.41 e 0.6 represents a meaningful concordance between the method result and the data [44]. When related to classification accuracy, it’s possible to enrich the classification analysis, considering the misclassifications error [45].

The benchmark permitted us to know the overfitting in the data, and a form of solving this issue was to use the Youden statistics. Based on the specificity and sensitivity during the validation process. The top four signatures had the coefficient of Youden in a range of 0.13 to 0.18 and were most proximal to 1 best in the classification [46], but the size of the study influenced this index [47].

The use of majority voting with the top four signatures results in the final signature of our work, composed of 7 mRNAs: (i) *PTTG1*, (ii) *BTBD11*, (iii) *HECW2*, (iv) *INSR*, (v) *RFLNB*, (vi) *HMMR*, (vii) *RASD1*, two lncRNAs: (i) *SNHG15*, (ii) *AF117829.1* and two miRNAs: (i) *hsa-miR-381-3p* e (ii) *hsa-miR-130a-3p*.

The validation with an external dataset is a process in the ML field to evaluate model generalization [48]. Our signature presents a great result, with accuracy and AUC of 72% and 81.5%, respectively. Other studies had constructed signatures associated with survival [31] and gene expression related to the immune system [49]in the literature.

### 4.2. Validation and Biological Interpretation

#### 4.2.1. Genomic and Functional Alterations

The somatic alterations of the coding genes from signature were more commonly associated with missense or frame_shit_del, except for the HMMR and RFLNB. Regarding the copy number variations, the amplified or deleted regions were not in the same location as the genes in the signature.

Analyzing the risk associated with survival or metastasis development showed a significant association of four genes from the gene signature. The lncRNA AF117829.1 and the miRNA hsa-miR-130a-3p were present in both analyses. The miRNA association is related to various cancers, such as bladder, breast, hepatocellular, glioma, and osteosarcoma [50–55]. Therefore, the presence of the PTTG1 and hsa.miR.130a.3p genes corroborate the literature, where in a situation of high expression is the poor prognosis, and for the hsa.miR.130a.3p and hsa.miR.381.3p are associated with metastatic development. However, the lncRNA remains unknown, and these features could be added to its actions, which are still under study.

The functional annotation resulted in very diversified pathways. The aldosterone-regulated sodium reabsorption pathway acts in sodium and potassium metabolism, and is a biomarker pathway for metastatic development and prognosis in ccRCC [56–58]. Another detected process was the longevity regulation pathway, which is affected by the caloric restriction related to mammalian feeding [59, 60] and regulates many other pathways, such as insulin signalization, PI3K/AKT, TNF, AMPK signalization, and mTOR pathway targets, that are also annotated as pathways regulated by the miRNAs targets. The TNF signaling pathway acts with PI3K/AKT and NF-kappa-B pathways for cellular necrosis, apoptosis, oncogenesis, and tumoral metastasis in many cancers [61].

The pathways related to the biological processes in both approaches, using the coding genes and miRNAs, showed annotation to cell cycle regulation, controlling the separation and segregation of sister chromatids, RNA polymerase II transcription, up-regulating and accommodating, its transcription activity of coding and non-coding genes [62], as well as processes related to cell-cell communication.

Thus, the functional annotation showed that the signature genes are associated with processes for metastatic development, associating them with relevant pathways such as PI3K/ATK and mTOR. When altered, these components trigger abnormal responses such as longevity and insulin regulation, all of them essential for cellular homeostasis

#### 4.2.2. Gene Signature Analysis in the ceRNA Network

As presented, the ceRNA network had a cluster distribution, showing dense regions with more presence of genes and fully connected, and sparse regions, with clusters more distant and without connectivity. A competition characteristic is observed in ten of the eleven genes in the signature. The processes related to the genes are the most diverse, like the regulation of cell motility by the HMMR [63], the regulation of the oncogenic pathways PI3K/AKT/mTOR by the *INSR* [64], the negative regulation of cell cycle by the PTTG1 and its action as an oncogene in the ccRCC microenvironment [65–67].

The lncRNA AF11782.1 mechanism of action remains unknown but was related to the proliferation, differentiation, and regulation of the immunity of T cells [68, 69], and as presented earlier, his expression was found to be related to metastatic development and worst prognosis of ccRCC patients, indicating new actions within the cancer field of studies. At this cluster, due to the high level of expression, a sponge act could be existing where the miRNA doesn’t degrade the POLE2 and HMMR, promoting cell differentiation and metastasis development.

The *RASD1* was the only gene that didn’t show a competition pattern and is responsible for the regulation of the RAS superfamily. In situations of increased expression, it is related to the reduction of the cell growth and the direction to apoptosis, acting in the opposite direction of the RAS family, associated with cell growth and tumor expansion [70]. As observed in his expression levels and his first ligands, the miRNA regulation is probably upon him, indicating the absence of the sponging action by the lncRNA, and promoting cancer cell proliferation.

## 5. Conclusion

This study aimed to build a transcriptional signature of clear cell renal cell carcinoma from differentially expressed genes that act as a Competitor Endogenous RNA network.

Using feature selection techniques for signature construction represents a promising application in this vast area of pattern recognition and machine learning. By integrating expression data with clinical information, we successfully constructed transcriptional signatures comprising multiple genes. The incorporation of evaluative metrics allowed us to gain valuable insights into the signature, assessing the metrics of accuracy, sensitivity, and specificity of the signature in order to classify metastatic tissue expression. Using the external dataset permitted the examination of the signature generalization, thus validating its action as a metastatic classifier in clear cell renal cancer.

With the cluster-by-cluster analysis, it was possible to know the actions performed by the signature genes within the cellular environment of clear cell renal cell carcinoma and how the effects of this regulatory process occur.

## Data availability statement

The study utilized openly accessible datasets for analysis. The findings presented in this paper stem from information gathered by the TCGA Research Network. The TCGA-KIRC dataset (version 07-19-2019) can be accessed through the UCSC Xena Browser[71], while the ICGC-RECA dataset is available via the ICGC Data Portal[27].

## Acknowledgments

The authors express their gratitude to Rafaella Ferraz and Iara de Souza for their valuable input and suggestions during the drafting of the manuscript. Additionally, the authors extend their thanks to the Multidisciplinary Bioinformatics Environment (BioME) at UFRN for generously providing the computing resources necessary for data processing.

## REFERENCES

1. Marcos Dall’Oglio, Miguel Srougi, Luciano Nesrallah. Câncer de Rim. In: Tratado de Clinica Médica. 2^a^. Rio de Janeiro; 2006. p. 3264–73.

2. Kumar V, Abbas AK, Fausto N, Robbins SL, Cotran RS. Robbins e Cotran: patologia: bases patológicas das doenças. 7. ed. Rio de Janeiro: Elsevier; 2008.

3. Muglia VF, Prando A. Renal cell carcinoma: histological classification and correlation with imaging findings. Radiol Bras. 2015;48:166–74.

4. NKF – National Kidney Fundation. Renal Carcinoma Guidelines. 2017.

5. Wang Y, Li Z, Li W, Zhou L, Jiang Y. Prognostic significance of long non-coding RNAs in clear cell renal cell carcinoma: A meta-analysis. Medicine (Baltimore). 2019;98:e17276.

6. Cui H, Shan H, Miao MZ, Jiang Z, Meng Y, Chen R, et al. Identification of the key genes and pathways involved in the tumorigenesis and prognosis of kidney renal clear cell carcinoma. Sci Rep. 2020;10:4271.

7. The Cancer Genome Atlas Research Network. Comprehensive molecular characterization of clear cell renal cell carcinoma. Nature. 2013;499:43–9.

8. Klinge CM. Non-coding RNAs: long non-coding RNAs and microRNAs in endocrine-related cancers. Endocr Relat Cancer. 2018;25:R259–82.

9. Kazimierczyk, Kasprowicz, Kasprzyk, Wrzesinski. Human Long Noncoding RNA Interactome: Detection, Characterization and Function. Int J Mol Sci. 2020;21:1027.

10. Statello L, Guo C-J, Chen L-L, Huarte M. Gene regulation by long non-coding RNAs and its biological functions. Nat Rev Mol Cell Biol. 2021;22:96–118.

11. Morris KV, Mattick JS. The rise of regulatory RNA. Nat Rev Genet. 2014;15:423–37.

12. Yao R-W, Wang Y, Chen L-L. Cellular functions of long noncoding RNAs. Nat Cell Biol. 2019;21:542–51.

13. Schmitz SU, Grote P, Herrmann BG. Mechanisms of long noncoding RNA function in development and disease. Cell Mol Life Sci. 2016;73:2491–509.

14. Wang P-S, Wang Z, Yang C. Dysregulations of long non-coding RNAs − The emerging “lnc” in environmental carcinogenesis. Semin Cancer Biol. 2021;76:163–72.

15. Chiu H-S, Somvanshi S, Patel E, Chen T-W, Singh VP, Zorman B, et al. Pan-Cancer Analysis of lncRNA Regulation Supports Their Targeting of Cancer Genes in Each Tumor Context. Cell Rep. 2018;23:297–312.e12.

16. Salmena L, Poliseno L, Tay Y, Kats L, Pandolfi PP. A ceRNA Hypothesis: The Rosetta Stone of a Hidden RNA Language? Cell. 2011;146:353–8.

17. Qi X, Lin Y, Chen J, Shen B. Decoding competing endogenous RNA networks for cancer biomarker discovery. Brief Bioinform. 2020;21:441–57.

18. Chan J, Tay Y. Noncoding RNA:RNA Regulatory Networks in Cancer. Int J Mol Sci. 2018;19:1310.

19. Bhan A, Soleimani M, Mandal SS. Long Noncoding RNA and Cancer: A New Paradigm. Cancer Res. 2017;77:3965–81.

20. Liu SJ, Dang HX, Lim DA, Feng FY, Maher CA. Long noncoding RNAs in cancer metastasis. Nat Rev Cancer. 2021;21:446–60.

21. Subramanian I, Verma S, Kumar S, Jere A, Anamika K. Multi-omics Data Integration, Interpretation, and Its Application. Bioinforma Biol Insights. 2020;14:117793221989905.

22. Reel PS, Reel S, Pearson E, Trucco E, Jefferson E. Using machine learning approaches for multi-omics data analysis: A review. Biotechnol Adv. 2021;49:107739.

23. Black JE, Kueper JK, Williamson TS. An introduction to machine learning for classification and prediction. Fam Pract. 2023;40:200–4.

24. Kann BH, Hosny A, Aerts HJWL. Artificial intelligence for clinical oncology. Cancer Cell. 2021;39:916–27.

25. Liu H, Motoda H, editors. Computational methods of feature selection. Boca Raton: Chapman & Hall/CRC; 2008.

26. Kuhn M, Johnson K. Feature engineering and selection: a practical approach for predictive models. Boca Raton London New York: CRC Press, Taylor & Francis Group; 2020.

27. Zhang J, Bajari R, Andric D, Gerthoffert F, Lepsa A, Nahal-Bose H, et al. The International Cancer Genome Consortium Data Portal. Nat Biotechnol. 2019;37:367–9.

28. Li R, Qu H, Wang S, Wei J, Zhang L, Ma R, et al. *GDCRNATools* : an R/Bioconductor package for integrative analysis of lncRNA, miRNA and mRNA data in GDC. Bioinformatics. 2018;34:2515–7.

29. Li J-H, Liu S, Zhou H, Qu L-H, Yang J-H. starBase v2.0: decoding miRNA-ceRNA, miRNA-ncRNA and protein–RNA interaction networks from large-scale CLIP-Seq data. Nucleic Acids Res. 2014;42:D92–7.

30. Shannon P, Markiel A, Ozier O, Baliga NS, Wang JT, Ramage D, et al. Cytoscape: a software environment for integrated models of biomolecular interaction networks. Genome Res. 2003;13:2498–504.

31. Terrematte P, Andrade D, Justino J, Stransky B, de Araújo D, Dória Neto A. A Novel Machine Learning 13-Gene Signature: Improving Risk Analysis and Survival Prediction for Clear Cell Renal Cell Carcinoma Patients. Cancers. 2022;14:2111.

32. Stawiski K, Kaszkowiak M, Mikulski D, Hogendorf P, Durczyński A, Strzelczyk J, et al. OmicSelector: automatic feature selection and deep learning modeling for omic experiments. preprint. Bioinformatics; 2022.

33. Lang M, Binder M, Richter J, Schratz P, Pfisterer F, Coors S, et al. mlr3: A modern object-oriented machine learning framework in R. J Open Source Softw. 2019;4:1903.

34. Mayakonda A, Lin D-C, Assenov Y, Plass C, Koeffler HP. Maftools: efficient and comprehensive analysis of somatic variants in cancer. Genome Res. 2018;28:1747–56.

35. Mermel CH, Schumacher SE, Hill B, Meyerson ML, Beroukhim R, Getz G. GISTIC2.0 facilitates sensitive and confident localization of the targets of focal somatic copy-number alteration in human cancers. Genome Biol. 2011;12:R41.

36. Therneau TM, Grambsch PM. Modeling survival data: extending the Cox model. 2. print. New York Berlin Heidelberg: Springer; 2001.

37. Harrison E, Drake T, Ots R. finalfit: Quickly Create Elegant Regression Results Tables and Plots when Modelling. R package version 1.0.6.

38. Aalen OO. A linear regression model for the analysis of life times. Stat Med. 1989;8:907–25.

39. Morris JA, Gardner MJ. Statistics in Medicine: Calculating confidence intervals for relative risks (odds ratios) and standardised ratios and rates. BMJ. 1988;296:1313–6.

40. Kanehisa M. Toward understanding the origin and evolution of cellular organisms. Protein Sci. 2019;28:1947–51.

41. The Gene Ontology Consortium, Carbon S, Douglass E, Good BM, Unni DR, Harris NL, et al. The Gene Ontology resource: enriching a GOld mine. Nucleic Acids Res. 2021;49:D325–34.

42. Wu T, Hu E, Xu S, Chen M, Guo P, Dai Z, et al. clusterProfiler 4.0: A universal enrichment tool for interpreting omics data. The Innovation. 2021;2:100141.

43. Vlachos IS, Zagganas K, Paraskevopoulou MD, Georgakilas G, Karagkouni D, Vergoulis T, et al. DIANA-miRPath v3.0: deciphering microRNA function with experimental support. Nucleic Acids Res. 2015;43:W460–6.

44. Landis JR, Koch GG. The measurement of observer agreement for categorical data. Biometrics. 1977;33:159–74.

45. Bendavid A. Comparison of classification accuracy using Cohen’s Weighted Kappa. Expert Syst Appl. 2008;34:825–32.

46. Youden WJ. Index for Rating Diagnostic Tests. 1950;3:32–5.

47. Zhou H. Statistical Inferences for the Youden Index. Atlanta, Geórgia.; 2011.

48. Ho SY, Phua K, Wong L, Bin Goh WW. Extensions of the External Validation for Checking Learned Model Interpretability and Generalizability. Patterns. 2020;1:100129.

49. Hua X, Chen J, Su Y, Liang C. Identification of an immune-related risk signature for predicting prognosis in clear cell renal cell carcinoma. Aging. 2020;12:2302–32.

50. Zhu J, Luo Y, Zhao Y, Kong Y, Zheng H, Li Y, et al. circEHBP1 promotes lymphangiogenesis and lymphatic metastasis of bladder cancer via miR-130a-3p/TGFβR1/VEGF-D signaling. Mol Ther. 2021;29:1838–52.

51. Chen J, Yan D, Wu W, Zhu J, Ye W, Shu Q. MicroRNA-130a promotes the metastasis and epithelial-mesenchymal transition of osteosarcoma by targeting PTEN. Oncol Rep. 2016;35:3285–92.

52. Li B, Huang P, Qiu J, Liao Y, Hong J, Yuan Y. MicroRNA-130a is down-regulated in hepatocellular carcinoma and associates with poor prognosis. Med Oncol. 2014;31:230.

53. Stückrath I, Rack B, Janni W, Jäger B, Pantel K, Schwarzenbach H. Aberrant plasma levels of circulating miR-16, miR-107, miR-130a and miR-146a are associated with lymph node metastasis and receptor status of breast cancer patients. Oncotarget. 2015;6:13387–401.

54. Ma F, Xie Y, Lei Y, Kuang Z, Liu X. The microRNA-130a-5p/RUNX2/STK32A network modulates tumor invasive and metastatic potential in non-small cell lung cancer. BMC Cancer. 2020;20:580.

55. Xu C-H, Xiao L-M, Liu Y, Chen L-K, Zheng S-Y, Zeng E-M, et al. The lncRNA HOXA11-AS promotes glioma cell growth and metastasis by targeting miR-130a-5p/HMGB2. Eur Rev Med Pharmacol Sci. 2019;23:241–52.

56. Connell JMC, Davies E. The new biology of aldosterone. J Endocrinol. 2005;186:1–20.

57. Wei W, Lv Y, Gan Z, Zhang Y, Han X, Xu Z. Identification of key genes involved in the metastasis of clear cell renal cell carcinoma. Oncol Lett. 2019. https://doi.org/10.3892/ol.2019.10130.

58. Zhang F, Wu P, Wang Y, Zhang M, Wang X, Wang T, et al. Identification of significant genes with prognostic influence in clear cell renal cell carcinoma via bioinformatics analysis. Transl Androl Urol. 2020;9:452–61.

59. Barzilai N, Huffman DM, Muzumdar RH, Bartke A. The Critical Role of Metabolic Pathways in Aging. Diabetes. 2012;61:1315–22.

60. Vara JÁF, Casado E, de Castro J, Cejas P, Belda-Iniesta C, González-Barón M. PI3K/Akt signalling pathway and cancer. Cancer Treat Rev. 2004;30:193–204.

61. Chu W-M. Tumor necrosis factor. Cancer Lett. 2013;328:222–5.

62. Schier AC, Taatjes DJ. Structure and mechanism of the RNA polymerase II transcription machinery. Genes Dev. 2020;34:465–88.

63. Hardwick C, Hoare K, Owens R, Hohn H, Hook M, Moore D, et al. Molecular cloning of a novel hyaluronan receptor that mediates tumor cell motility. J Cell Biol. 1992;117:1343–50.

64. Takahashi M, Inoue T, Huang M, Numakura K, Tsuruta H, Saito M, et al. Inverse relationship between insulin receptor expression and progression in renal cell carcinoma. Oncol Rep. 2017;37:2929–41.

65. Sun Y, Liu W-Z, Liu T, Feng X, Yang N, Zhou H-F. Signaling pathway of MAPK/ERK in cell proliferation, differentiation, migration, senescence and apoptosis. J Recept Signal Transduct Res. 2015;35:600–4.

66. Mei L. Multiple types of noncoding RNA are involved in potential modulation of PTTG1’s expression and function in breast cancer. Genomics. 2022;114:110352.

67. Zi Z. Molecular Engineering of the TGF-β Signaling Pathway. J Mol Biol. 2019;431:2644–54.

68. Xia F, Yan Y, Shen C. A Prognostic Pyroptosis-Related lncRNAs Risk Model Correlates With the Immune Microenvironment in Colon Adenocarcinoma. Front Cell Dev Biol. 2021;9:811734.

69. Li Y, Deng L, Pan X, Liu C, Fu R. The Role of lncRNA AF117829.1 in the Immunological Pathogenesis of Severe Aplastic Anaemia. Oxid Med Cell Longev. 2021;2021:1–19.

70. Vaidyanathan G, Cismowski MJ, Wang G, Vincent TS, Brown KD, Lanier SM. The Ras-related protein AGS1/RASD1 suppresses cell growth. Oncogene. 2004;23:5858–63.

71. Goldman MJ, Craft B, Hastie M, Repečka K, McDade F, Kamath A, et al. Visualizing and interpreting cancer genomics data via the Xena platform. Nat Biotechnol. 2020;38:675–8.

