## Supplemental Figure 1 for "Machine Learning Gene Signature to Metastatic ccRCC based on ceRNA Network"

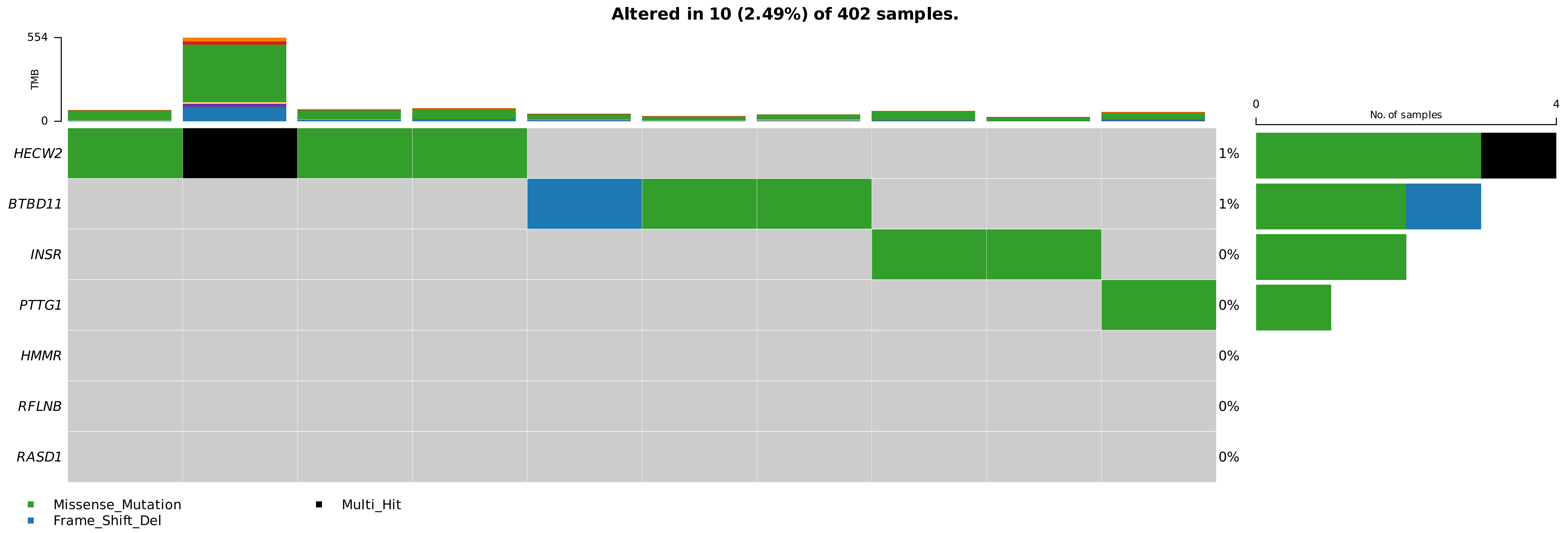


**Figure S1:** Oncoplot with the mutations recorded in the signature coding genes, The bar graph on the right representes the amount of mutated samples and the bar graph above representes the mutations registeres on these samples

**A**

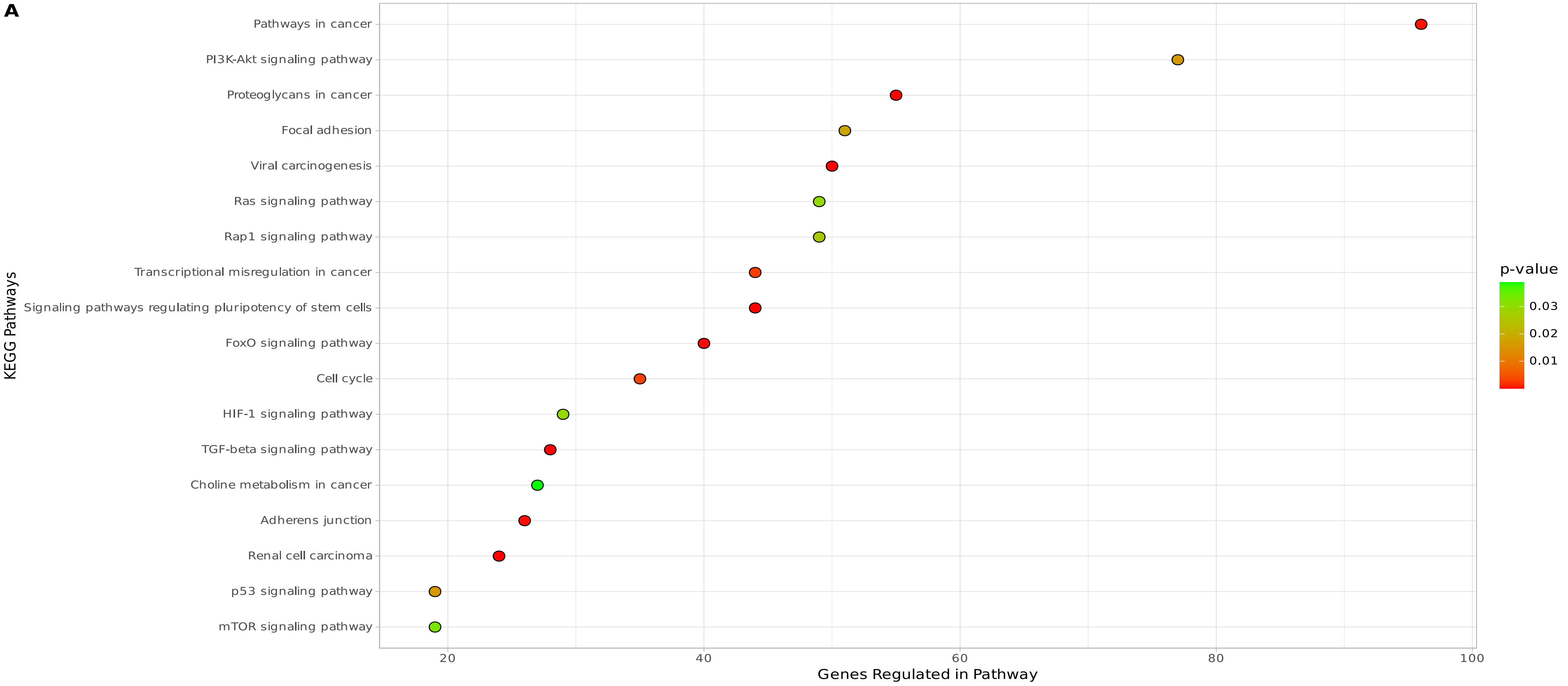


**B**

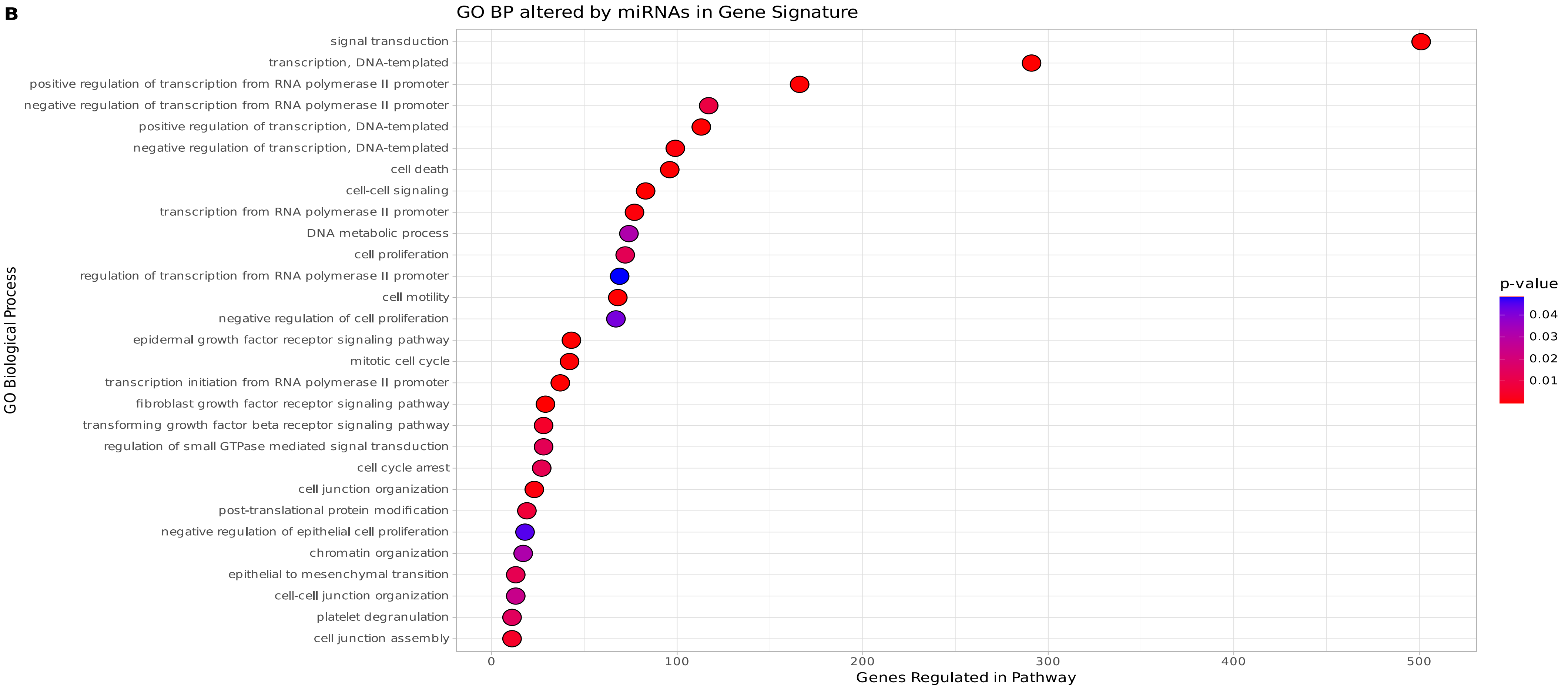


**FIgure S2:** Functional annotation made from (a) KEGG and (b) Gene Ontology using the targets of miRNAs participating in the signature. In both, the Y axis represents the annotated pathways in the databases and the X axis represents the number of genes regulated by the miRNAs in the pathway.

Table S1: Feature selection techniques and application stage.

| Method | Technique | Step |
|---|---|---|
| Filter | Generalized Linear Model | Feature Selection and Benchmarking |
| | Linear Discriminant Analysis | Feature Selection |
| | Akaike Information Criterion | Feature Selection |
| | eXtreme Gradient Boosting | Benchmarking and Validation |
| Wrapper | Boruta | Feature Selection |
| | Recursive Feature Elimination | Feature Selection |
| | Lasso | Feature Selection |
| | ElasticNet | Feature Selection |
| | Support Vector Machine | Benchmarking and Validation |
| | Naive Bayes | Validation |
| | k-Nearest Neighbors | Validation |
| | Random Forest | Benchmarking and Validation |
